# Ethanol tolerance in *Escherichia coli* DH5-Alpha developed by serial exposure to sublethal doses is conferred to wild strains by horizontal gene transfer

**DOI:** 10.1101/2020.01.20.912196

**Authors:** Ronak Borana, Shreya Bari

## Abstract

Microorganisms evolve novel mechanisms and pathways to mitigate various stresses. These are developed by the accumulation of beneficial mutations over many generations. Such adaptations are often transferred from mutant microorganisms that develop it intrinsically to wild ones through horizontal gene transfer (HGT). It allows the latter to acquire favourable traits without having to employ resources to natively evolve. We ascertained this in *Escherichia coli* by first developing tolerance to ethanol, a potent disinfectant, by laboratory evolution and then transmitting it to the wild strain by HGT. Naturally, wild type *E. coli* cannot survive beyond 35% v/v ethanol in LB media. By serially increasing the concentration of ethanol by 5% v/v and selecting the surviving colonies, we were able to impose an artificial selection pressure. This in vitro microevolution increased the ethanol tolerance in our mutant to 75% v/v. To test if this tolerance could be transferred to the wild strain through HGT, we meticulously exposed the ancestral wild *E. coli* to the newly tolerant mutants to facilitate the exchange of genetic material. After our exposure, the unadapted wild type *E. coli* acquired tolerance up to 55% v/v through transformation and 45% v/v through transduction. The baseline tolerance of 35% v/v remained unchanged after conjugation. While our results are still preliminary, they provide interesting insights into the role horizontal gene transfer in developing resistance to bacteriocidal stressors.

## Introduction

Adaptation is significant not only for neutralizing stress but also for ameliorating genetic diversity and speciation. It is a paramount force that drives evolution. It is studied by observing a test organism adapt to choreographed stresses under a controlled environment. This method is popularly called a laboratory or experimental evolution. It is conducted on many model organisms ranging from fruit flies to foxes. Along with testing evolutionary hypotheses, laboratory evolution is also used in developing vaccines^[1]^ and studying antibiotic resistance. At its core, it is exposing a test organism to a novel intervention for many generations and then comparing it with the ancestral population which essentially become the control arm. Microorganisms are an apt candidate for laboratory evolution because of their rapid generation time, easy handling and widely studied and accessible genomes. Sublethal doses of bacteriocidal agents like antibiotics cause sufficient bottlenecks in microbial populations that enforce the survival of only those organisms that adapt. Under controlled laboratory environment, this adaptation can happen in just a few hundred generations. Most laboratory evolution in microbes is usually conducted in nosocomial pathogens or model organisms like *E. coli*. The most noted example is of Lenski et al’s Long Term Experimental Evolution (LTEE) which has been studying laboratory evolution in *E. coli* for more than 30 years and 60,000 generations ^[2]^.

Laboratory evolution has been used in developing resistance to a range of different bacteriocidal agents. In a recent 2018 study^[3]^, *Vanco*mycin-resistant enterococci (VRE) were serially exposed to chlorohexidine, a gluconate disinfectant, frequently used in hospitals. In 21 days, the VRE developed a 4 fold tolerance to the initial MIC of chlorohexidine. Similarly, laboratory evolution has been used to generating tolerance to a range of biocides like saline stress^[4]^, antimicrobial peptide^[5]^, and aminoglycosides like kanamycin^[6]^ among others.

Ethanolgenic *E. coli* lacks the ability to naturally tolerate ethanol in its environment. This significantly hinders its ability to ferment ethanol, a promising biofuel, once the product starts to accumulate in the surrounding. Using laboratory evolution, a two-fold increase in tolerance was developed artificially in about 350 generations^[7]^. A natural example of this adaptive evolution to alcohol was noted in large 2018 study that found nosocomial *Enterococci* were becoming resistant to alcohol-based disinfectants^[8]^. The study compared isolates of *Enterococcus faecium* collected from hospitals between 1997-2015. The newer samples were significantly more tolerant of alcohol-based hand rubs than the older ones. They also found that alcohol tolerant VRE infected mice more quickly than their non-tolerant strain. While not yet being a threat to hospital hygiene management^[9] [10]^, pathogens tolerating higher concentrations of disinfecting alcohol is an alarming discovery. We decided to study this tolerance to alcohol, ethanol in our case, by serially exposing non-pathogenic *E. coli* to serially increasing sublethal concentrations of ethanol.

Horizontal or lateral gene transfer (HGT) is the exchange of any genetic material (genomic DNA/RNA, extrachromosomal structures etc) from one organism to another by any means except, reproductive or parent to offspring transfer which is the vertical transfer of gene. HGT is a powerful force in evolution as it helps the receiving organism acquire traits without spending the resources to naturally adapt to it. While most HGT’s have a deleterious or neutral effect some of them often confer an evolutionary advantage. Release, uptake and integration of exogenous genes is a highly variable process and has a very poor success rate. Despite this, when it works, it significantly increased the evolutionary fitness of the recipient. Some genes that have been acquired through HGT are responsible for conferring virulence, resistance to antibiotics, pesticide degrading ability, extreme pH tolerance among others.

HGT is observed not only between different species of bacteria but also between different domains of like like archaea and algae^[10]^. The similarity between sequences suggests^[11]^ that many eukaryotic genes owe their existence to HGT. They were most likely shuttled by viruses through transfection.

HGT operates through a wide range of mechanisms. Transformation, conjugation and transduction are the most known ones. Transformation occurs when a recipient microorganism accepts foreign genetic material through its cell membrane from the environment and incorporates it into its own genome. The uptaken genetic material can either be the genomic DNA or plasmid. The latter, however, is less efficient^[12].^ Transduction is a transfer of genetic material, mostly parts of genomic DNA from one microorganism to another by a virus or a phage. While most transductions are fatal for the recipient cell, bacteriophages are known to suttle important traits like antibiotic resistance across certain bacterial genera^[13].^ Conjugation, often associated with fertility, is a plasmid-mediated transfer of genes that occurs during cell to cell contact via an intercellular bridge called a pilus. Antibiotic conferring plasmid has been transmitted between different bacterial species using conjugative transfer^[14]^. There are other mechanisms like conjugative transposons that facilitated HGT but are relatively understudied ^[15]^.

Computation analysis of phylogeny shows that a large number of ancestral bacterial genes have been originally acquired via horizontal gene transfer^[16][17]^. It also plays a paramount role in antimicrobial resistance. We decided to develop in vivo artificial tolerance to ethanol so that we could study the impact and the scope of HGT in exchanging novel mutations that incapacitate bacteriocidal action.

## Methods and Materials

### Media and Strains

DH5-Alpha *Escherichia coli* was selected as the test organism. It is an artificially engineered strain of the model organism *E. coli* which has increased transformation efficiency and well-defined protocols maintenance and competency induction. This trait is used to introduce ampicillin resistance plasmid as a distinguishing marker later in the experiment. Lysogeny Broth or LB media was used for all the serial transfers. Super optimal broth or SOB which is slightly more favourable^[18]^ than LB was used for HGT experiments. Broth with varied ethanol concentration was made by suing ethanol to make up the volume of LB broth after sterilization it. For instance, to make 40% v/v, 2.5 gm of solid LB composition was dissolved in 60 ml distilled water. After autoclaving it, 40 ml of 100% ethanol was added to make the final volume of the broth to 100 ml.

### Serial Transfer for Laboratory evolution

The higher concentrations of ethanol in LB media (+50% v/v) were very cloudy and the optical density readings of those broths were not reliable or reproducible. Therefore CFU/ml were used to as a proxy for growth. Because of the strong bactericidal effect of ethanol, the growth was severely inhibited at higher concentrations (+55% v/v). To remedy this, 0.5 ml was spread on LB plates instead of 0.1ml and the number of viable colonies was multiplied by 2 to obtain the final CFU/ml reading.

Lyophilized *E. coli* DH5 Alpha cells were revived in 100 ml LB broth. This was the ancestral wild type population that was used as the seed culture for all experiments. The natural or baseline ethanol tolerance was calculated by inoculating 1 ml of the revived *E. coli* in +5% v/v increasing concentration of ethanol-LB broth from 5-100% v/v. CFU/ml was measured after 24 hours of incubation on a rotatory shaker. 40% v/v was the MIC and the CFU/ml reading of 35% v/v was 28. Broths with +40% v/v were allowed to incubate for more 24 hours after which the CFU readings were taken. This was continued till the CFU/ml reading of all concentrations above 35% v/v were 0 for 8 days. Thus the highest tolerance for the wild unadapted population was estimated to be 35% v/v ethanol in LB broth.

For serial transfer, 0.5 ml of revived *E. coli* was added to a 50 ml 30% v/v ethanol-LB broth. After 24 hours of incubation on a rotatory shaker, 0.5ml of the 30% v/v broth was added to 35% v/v. The CFU/ml of this 35% v/v was taken every 24 hours until a non-zero CFU reading was obtained. When there was a non zero reading, it was assumed that tolerance has been developed and 0.5 ml from 35% v/v was then inoculated in 40% v/v. This process was serial transfer was repeated for every +5% v/v until there were was no non-zero CFU/ml reading for 8 consecutive days or 192 hours of incubation. CFU and OD readings helped in ensuring that the test organism is challenged with relatively higher ethanol concentration (+5% v/v) only when there were viable cells in the lower concentration. The number of days it took to develop tolerance, the first non zero CFU and the total difference in OD at 540 nm was plotted for all the concentrations where growth was recorded.

The bacteriocidal nature of ethanol substantially decreased the likelihood of any contamination. Yet to ensure that all transfers were aseptic, a differential selective agar and staining was used to ensure fidelity.

### Horizontal Gene Transfer Experiments

Experiments involving the transfer and modification of genetic materials like horizontal gene transfer require access to sequencing and advanced bioinformatics expertise. Therefore we confided only to basic preliminary molecular biology experiments.

To differentiate ancestral unadapted test bacteria (wild strain) from newly tolerant ones (mutant strain) ampicillin-resistant plasmid were used as differential markers. Hanahan’s protocol 18 for transformation was used to induce competence in the wild strain. Plasmids with the ampicillin-resistant gene were transformed into the wild strain to make them resistant to ampicillin. For all further experiments, the wild strain was now resistant to amp+ media whereas the mutant strain was not. This allowed in differential selection.

Ethanol tolerance was measured after the experiments by inoculating 0.5 ml of the test in 30, 35, 40, 45….80% v/v ethanol-LB broth. Growth in these concentrations was established by OD difference and CFU/ ml readings every 24 hours for 8 days until a non-zero reading of CFU/ml was obtained. Similar to how the initial baseline tolerance of 35% v/v was established in the wild strain.

To transfer the mutated genetic material that granted tolerance to 75% v/v ethanol by transformation, Hanahan’s protocol 18 was used. Cells from the mutant strain were lysed in a DNase and RNase free medium and their genetic material were isolated by centrifugation. Competent cells from the wild strain were transformed with this genetic material isolated from the mutants. After this, the transformed bacteria were grown in amp+ media to ensure that they are not contaminated with the mutant cells. The ethanol tolerance of the wild strain that had not been exposed to ethanol before, after transformation was measured. It was then compared with a control which was made cells from the wild strain that were not transformed.

For transduction, viruses from the culture containing the mutant strain were isolated using phage titration. They were transferred to amp+ media containing the cells from the wild strain. The ampicillin in the media ensured that the growth of any mutant cells in the phage titrate would be inhibited. The phages that earlier infected the mutants could incorporate some of the host’s genes in their genome and subsequently transfer them to the wild cells when they are allowed to infect them. The ethanol tolerance in the transduced wild cells was measured and compared with appropriate control.

Similarly, conjugation was carried out by co-growing the wild and the mutant strains together and allowing only the wild cells to survive by adding ampicillin in the culture. The surviving wild type cells were isolated and their ethanol tolerance was measured.

## Results

### Serial Transfer for Laboratory evolution

Short-chain alcohols like isopropanol and ethanol are strong sterilizing agents and are regularly used in laboratories and hospitals. Along with being bacteriocidal, alcohols are also fungicidal and virucidal and tuberculocidal. The mode of action of alcohol is known to be by penetrating the hydrocarbon component^[19]^ of the phospholipid bilayer and causing disorganization and loss of integrity of the membrane causing the discharge of the intracellular components. Some studies also note that the bacteriocidal nature of alcohol is likely due to its ability to coagulate and denature the transmembrane proteins^[20]^. Ethanol particularly also shows some secondary effects on *E. coli* like inhibition of synthesis of proteins, peptidoglycan and destruction of essential enzymes like dehydrogenase^[21]^.

Bacteria can evolve tolerance to ethanol by many different mechanisms. They can do so by metabolizing ethanol and utilizing it as a signalling molecule^[22]^. *Acinetobacter baumannii* a notorious nosocomial pathogen, that is responsible for hospital-acquired Multidrug-resistant (MDR) Pneumonia is getting more tolerant to alcohol-based hand rubs. It is doing so by secreting a protein called OmpA^[23]^. *E. coli* also secretes OmpA^[24]^. Strengthening of the opposing potassium and proton electrochemical membrane gradient is also known to increase alcohol tolerance^[25]^.

A 2010 study^[26]^ in *E. coli* used whole-genome fitness profiling to pin ethanol tolerance to rewiring of regulatory and metabolic pathways. They found that perturbation of several genes associated with modulation of the acid stress response, osmotolerance, cell wall biosynthesis, assimilation and degradation of ethanol in the TCA cycle were responsible for the increased tolerance. Therefore tolerance to ethanol can be generated through a combination of many such altered metabolic mechanisms.

In our laboratory evolution of *E. coli* DH5 Alpha, we were able to make the ancestral wild strain which had a baseline ethanol tolerance of 35% v/v more tolerant by up to 75% v/v. We were able to achieve this over a period of 46 days of serial transferring. 0.5ml of 75% v/v tolerant mutant were transferred to 80% v/v. After 8 days of incubation, there still were no viable cells in the 80% v/v. We inferred 80% v/v to be the MIC for the mutant and concluded the serial transfer. Overall, there was a 114% increase in ethanol tolerance after the laboratory microevolution.

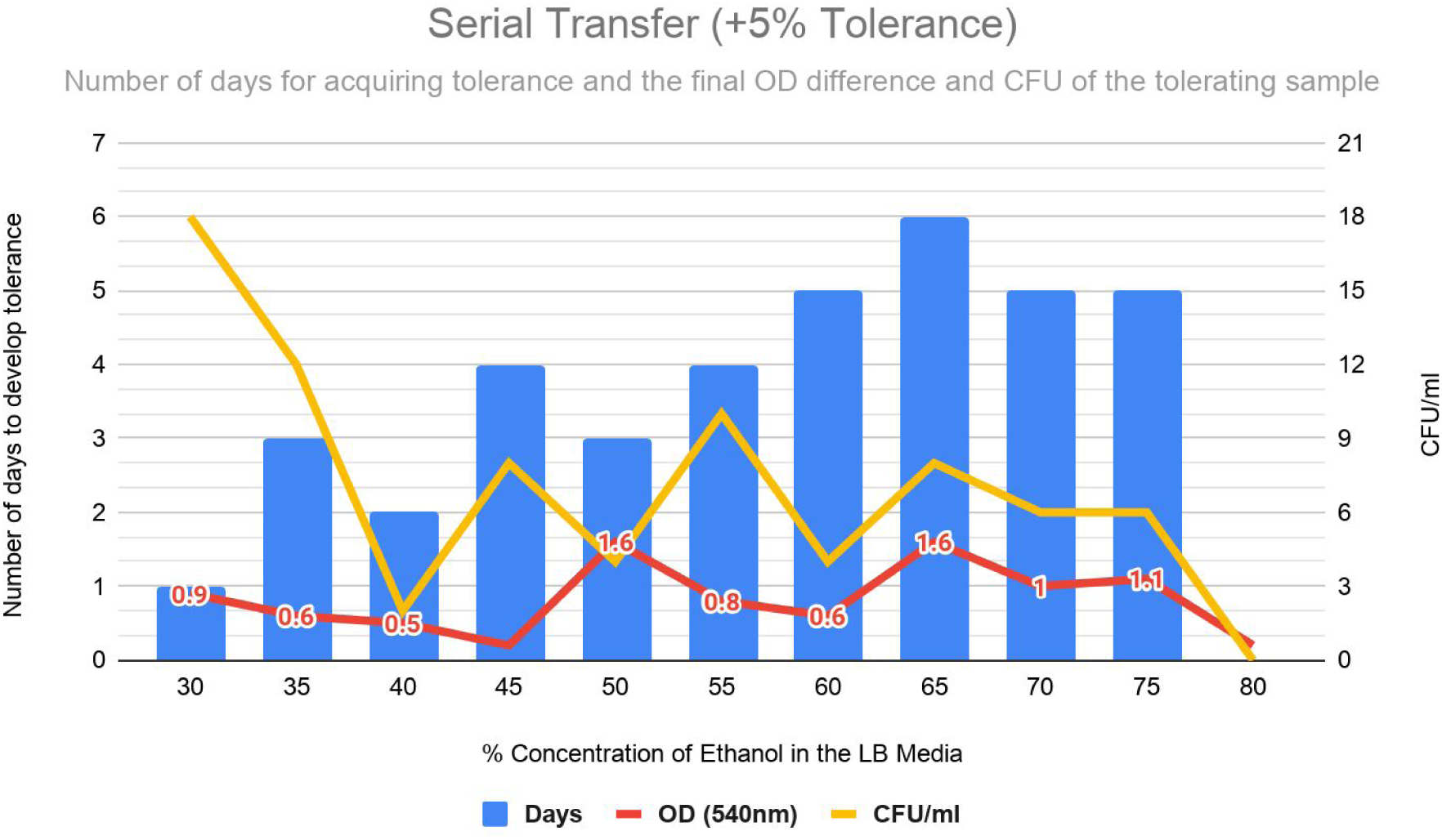

### Horizontal Gene Transfer Experiments

An increase in ethanol tolerance was seen after transformation and transduction but not in conjugation. As compared to the unadapted wild bacteria that had a maximum tolerance of 35% v/v (control), the transformed bacteria had a maximum tolerance of 55% v/v which is a 57 per cent increase. Wild strain after transduction intervention too had a 28 per cent increase in maximum tolerance at 45% v/v. The wild cells after conjugation, however, showed no increase in ethanol tolerance and was still capped at 35% v/v.

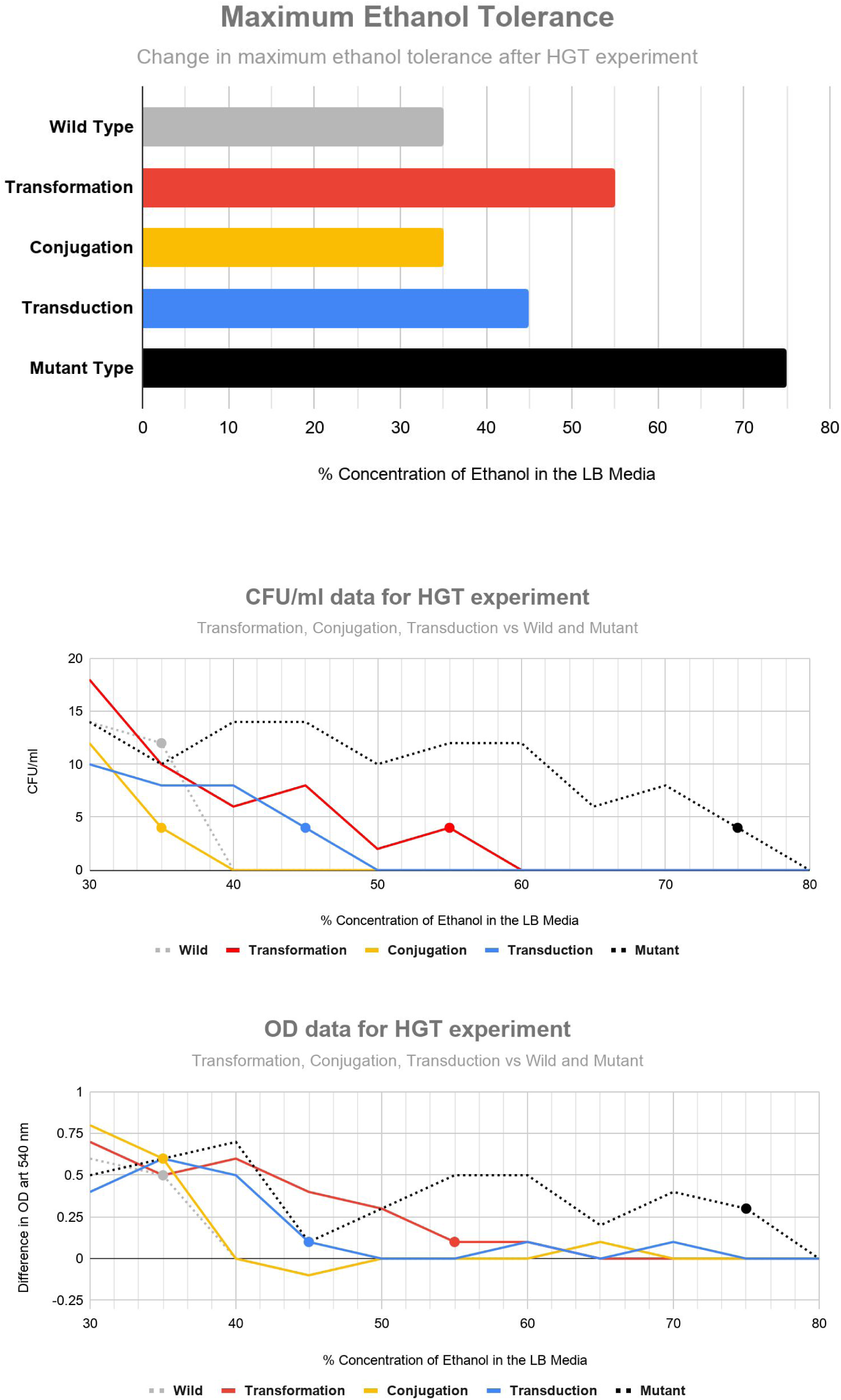

## Discussions

The results from our HGT experiment establish that ethanol tolerance as a trait can be conferred to unadapted wild strain by means of horizontal gene transfer. Transformation and transduction to be specific. The 114% gradual increase in ethanol tolerance is very surprising. It attests the significance of laboratory evolution as a powerful method to study adaptation in particular and evolutionary biology in general. The experiments we used to test HGT were very preliminary and therefore despite being positive, they are not completely conclusive. Further research is required to study the exact mechanism of tolerance, the mutations that enable them and the rate and the process of exchange of these mutations through horizontal gene transfer. All the intermediary mutants in our experiment have been carefully preserved in agar slants for prospective future studies on the above-mentioned areas.

Understanding adaptation is not only important for the advancement of evolutionary biology but also addressing the growing crisis of resistance to antimicrobials like antibiotics^[27]^ and alcohol-based sanitizer^[8]^. Such artificial in vivo microevolution can provide valuable insights into the mechanism of such resistance^[28]^ and provide a controlled setting to test our mitigating strategies. Along with antimicrobial resistance, understanding the stress adaptation can also help in developing more efficient strains ^[29] [30]^ to optimize the fermentation of commercial drugs, biologics and other therapeutic metabolites.

